# Proteomics analysis reveals a role for *E. coli* polyphosphate kinase in membrane structure and polymyxin resistance during starvation

**DOI:** 10.1101/2023.07.06.546892

**Authors:** Kanchi Baijal, Iryna Abramchuk, Carmen M. Herrera, M. Stephen Trent, Mathieu Lavallée-Adam, Michael Downey

## Abstract

Polyphosphates (polyP) are chains of inorganic phosphates that can reach over 1000 residues in length. In *Escherichia coli*, polyP is produced by the polyP kinase (PPK) and is thought to play a protective role during the response to cellular stress. However, the molecular pathways impacted by PPK activity and polyP accumulation remain poorly characterized. In this work we used label-free mass spectrometry to study the response of bacteria that cannot produce polyP (Δ*ppk*) during starvation to identify novel pathways regulated by PPK.

In response to starvation, we found 92 proteins significantly differentially expressed between wild-type and Δ*ppk* mutant cells. Wild-type cells were enriched for proteins related to amino acid biosynthesis and transport, while *Δppk* mutants were enriched for proteins related to translation and ribosome biogenesis, suggesting that without PPK, cells remain inappropriately primed for growth even in the absence of required building blocks.

From our dataset, we were particularly interested in Arn and EptA proteins, which were downregulated in Δ*ppk* mutants compared to wild-type controls, because they play a role in lipid A modifications linked to polymyxin resistance. Using western blotting, we confirm differential expression of these and related proteins, and provide evidence that this mis-regulation in Δ*ppk* cells stems from a failure to induce the BasS/BasR two-component system during starvation. We also show that Δ*ppk* mutants unable to upregulate Arn and EptA expression lack the respective L-Ara4N and pEtN modifications on lipid A. In line with this observation, loss of *ppk* restores polymyxin sensitivity in resistant strains carrying a constitutively active *basR* allele.

Overall, we show a new role for PPK in lipid A modification during starvation and provide a rationale for targeting PPK to sensitize bacteria towards polymyxin treatment. We further anticipate that our proteomics work will provide an important resource for researchers interested in the diverse pathways impacted by PPK.

## INTRODUCTION

Polyphosphates (polyP) are homopolymers of inorganic phosphates joined together by high energy phosphoanhydride bonds. Although polyP is found across diverse organisms from bacteria to humans, the intracellular concentrations, and mechanisms by which it is produced vary widely [1–3]. In *Escherichia coli* (*E. coli*), polyP is synthesized by the polyphosphate kinase PPK and degraded by the exopolyphosphatase PPX [4]. In general, *E. coli* produce little to no detectable polyP when undergoing logarithmic growth in nutrient rich media [5]. However, in response to diverse cellular stressors including oxidative stress caused by exposure to hypochlorous acid (bleach) [6], heat shock [7], and nutrient starvation [8], PPK rapidly synthesizes polyP using ATP or GTP as a co-substrate [1]. This stress-induced population of polyP has been linked to protein folding and turnover [6, 7], transcriptional [9, 10] and translational control [11], and the regulation of bacterial heterochromatin [12]. In some cases, polyP is thought to impart these changes by interacting directly with protein targets to modulate their activity. For example, polyP produced during nutrient downshift interacts with the Lon protease to direct its activity towards degradation of ADP-bound DnaA and ribosomal proteins [13–15]. Collectively, these pathways inhibit DNA replication, while increasing the intracellular pool of amino acids to help *E. coli* adapt to changing conditions [13, 15]. PolyP can also function by chelating cations, for example as an inhibitor of the Fenton reaction in which iron catalyzes the formation of reactive oxygen species [16]. *E. coli ppk* mutants (e.g. Δ*ppk*) display increased sensitivity to cellular stress [6, 7, 17] and decreased motility [18], biofilm formation [17], and virulence [19–21]. While the molecular events underlying these phenotypes are not always known, the role of PPK enzymes as regulators of survival during cell stress is conserved across the bacterial kingdom. Notably, in addition to polyP, PPK enzymes also show enzymatic activity against nucleoside diphosphates [22–24], although the degree to which these functions contribute to stress resistance is unclear.

The PPK status of pathogenic *E. Coli* is an important regulator of infectivity in mouse models of infection [19, 25]. It has been suggested that polyP released by *E. coli* may play an important role in the reprogramming of macrophages, and this may involve polyP interaction with host receptors on the cell membrane or entry into host cells [19]. PPK has also been proposed as a novel target for various bacterial infections [17, 26]. It is noteworthy that mesalamine, a drug used to treat ulcerative colitis and Crohn’s disease, inhibits PPK enzymes *in vitro* and can reduce infection in a murine model in a PPK-dependent manner [17]. The pursuit of PPK as a valid therapeutic target demands a thorough understanding of how PPK impacts bacterial stress response at a systems-wide level.

To better understand the role of PPK and polyP in bacterial stress responses, we used label-free proteomics to identify proteins up- or downregulated in Δ*ppk* mutant cells relative to wild-type controls undergoing prolonged starvation, when polyP levels are high. We report that mutant Δ*ppk* cells fail to upregulate pathways required for amino acid biosynthesis and instead are enriched for processes related to ribosome biogenesis. In follow up work, we show a role for PPK in the modification of lipid A – the lipid anchor of the lipopolysaccharide (LPS) at the cell surface of Gram-negative bacteria [27]. We demonstrate that during starvation, PPK is required for expression of EptA and the Arn proteins, and their respective phosphoethanolamine (pEtN) and 4-amino-4-deoxy-L-arabinose (L-Ara4N) lipid A modifications, as well as for expression of the upstream BasS-BasR two-component system. In an antibiotic-resistant strain background, cells lacking *ppk* display increased susceptibility to the cationic antimicrobial peptide polymyxin B. Together, our work provides a novel resource for investigating molecular functions of polyP and new insights into how PPK inhibition could be best exploited in the clinic.

## RESULTS

### Proteomic differences between wild-type controls and Δ*ppk E. coli* upon nutrient starvation

We used label-free mass spectrometry analysis to compare proteomic differences between wild-type and Δ*ppk* cells following a shift from LB to MOPS minimal media (**Fig. 1A**). In the bacterial polyP field, a shift from nutrient rich to MOPS minimal media is commonly used to trigger polyP accumulation [5, 28, 29]. At the 3-hour time point used for analysis, all five replicates of wild-type cells showed accumulation of polyP, whereas Δ*ppk* mutants did not (**Supplementary Fig. 1 A**). Bioinformatics analysis of mass spectrometry data uncovered 1909 proteins total, of which 78 were significantly differentially expressed between the two conditions (FDR-adjusted *p*-value < 0.05) (**Fig. 1B, Supplementary Table 1**). In addition, 14 proteins were classified as all-or-none (detected in 0 replicates of Δ*ppk* mutant cells, but detected in all 5 replicates of wild-type cells, or vice versa) (**Figs. 1C**). We used western blotting to confirm expression differences for 6 (ArnB, ArnC, MetE, YbdL, YeaG, OtsA) out of the 7 top hits, validating the overall high-quality of the data set (**Fig. 1D**). Only RaiA failed to confirm in targeted western blotting experiments, showing inconsistent results between replicates.

**Figure 1.**
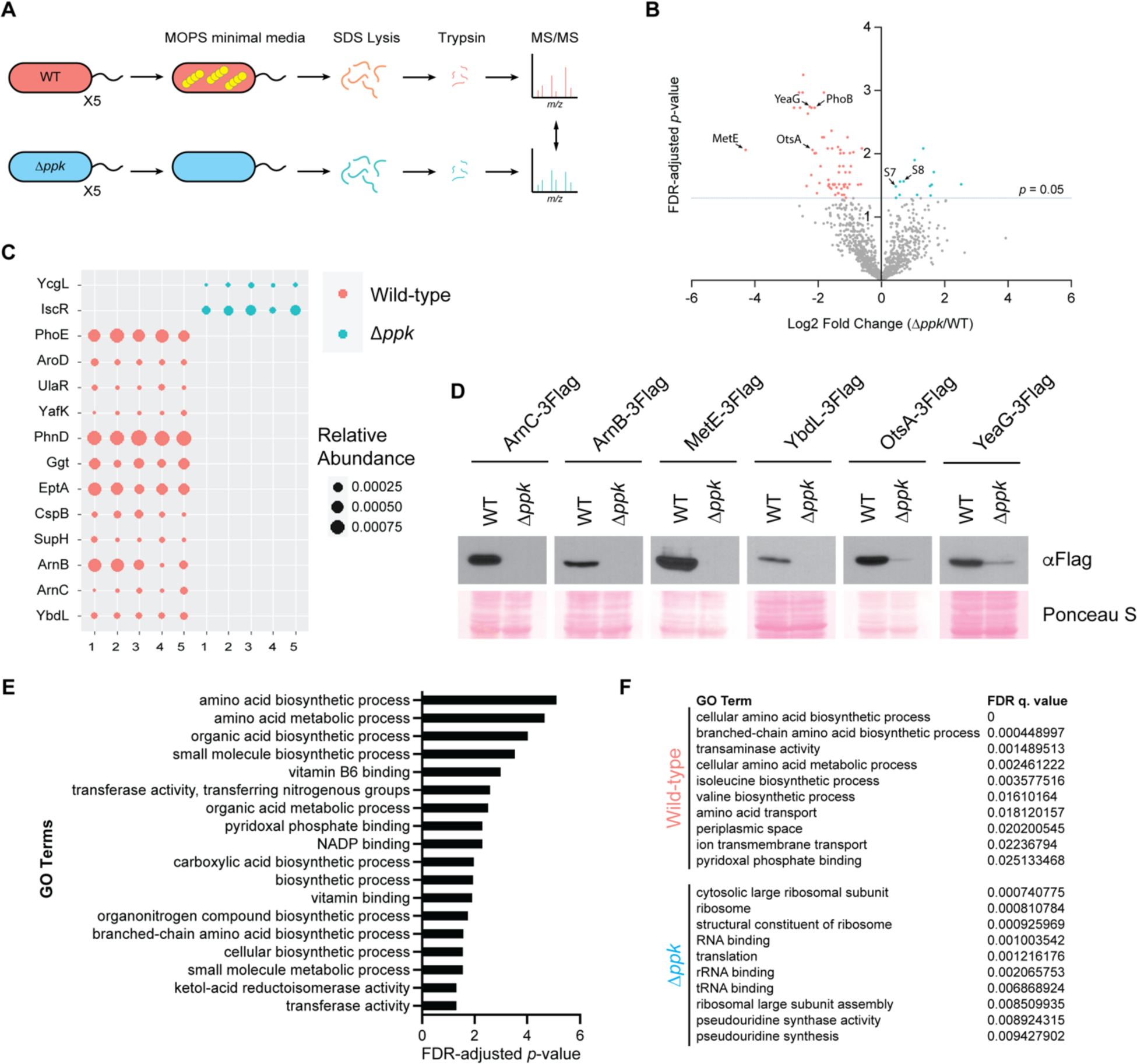
Broad proteomic changes in Δ*ppk* cells during stress. **A)** Experimental set up for proteomics analysis. Cells were grown in LB media to mid-exponential phase before a shift into MOPS minimal media (0.1 mM K_2_HPO_4_, 0.4% glucose) for 3 hours to induce amino acid starvation and polyP accumulation. The experiment was conducted using n=5 biological replicates. **B)** Volcano plot of significantly differentially expressed proteins (log2(fold-change Δ*ppk/*WT)). In red and blue are the significantly upregulated proteins (FDR-adjusted *p*-value < 0.05) in wild-type and Δ*ppk* strains, respectively. **C)** Bubble plot showing the ‘all-or-none’ proteins detected only in either wild-type cells or Δ*ppk* mutants. Data represent the normalized protein spectral counts in 5 biological replicates from each condition. **D)** Select confirmations of mass spectrometry data. Endogenously C-terminal 3Flag-tagged strains were grown under the same conditions for the mass spectrometry analysis. Protein extracts were resolved using a 12% (for YbdL) and 10% (for all other proteins) SDS-PAGE gel, transferred to PVDF membrane, and probed using an anti-Flag antibody. Images are representative of results from ≥3 experiments. **E)** GO terms that are significantly enriched among the differentially expressed and ‘all-or-none proteins’ identified by mass spectrometry analysis. **F**) GO terms deemed differentially expressed based on GSEA for wild-type and Δ*ppk* mutant cells.

Previous work by Varas *et al.* used mass spectrometry to compare proteomes of wild-type controls and Δ*ppk* mutants in nutrient rich LB media [30], where there is no detectable polyP accumulation [5, 31]. There, the authors identified 60 proteins upregulated and 32 proteins downregulated in Δ*ppk* cells [32]. The overlap between these differentially expressed proteins and the dataset described in our work is poor **(Supplementary Table 2)**. This suggests that there are vast proteomic differences between bacteria experiencing stress compared to those grown in LB media and most importantly, that there are unique roles of PPK and polyP in proteomic regulation during starvation that are uncovered by our work.

We performed Gene Ontology (GO) [33] enrichment analysis on the significantly differentially expressed and all-or-none proteins (92 total) and identified 16 enriched GO terms (FDR-adjusted *p*-value < 0.05) (**Fig. 1 E**). These included terms related to amino acid, organic acid, and small molecule biosynthesis. We next used Gene Set Enrichment Analysis (GSEA) [34] on the entire data set of 1909 proteins to look for GO terms that are differentially expressed between the wild-type cells and Δ*ppk* mutants. This analysis showed that wild-type cells were enriched for proteins related to amino acid biosynthesis and transport (**Fig. 1F**). In contrast, Δ*ppk* mutant cells were enriched for proteins broadly related to ribosome biogenesis and translation (**Fig. 1F**). These data point to a model wherein Δ*ppk* mutants fail to properly respond to starvation; the cells remain primed for growth while failing to activate pathways to increase the availability of amino acids and other biomolecules needed for that purpose. In line with this interpretation, polyP interacts with the Lon protease to promote degradation of ribosomal proteins including S2, L9 and L13, as well as nucleoid proteins such as HupA, HimA (IhfA) and translational elongational protein InfC [13, 35]. This degradation has been proposed to provide free amino acids to allow for targeted translation during starvation [13]. In agreement with this data, we detected significant upregulation of 30S ribosomal proteins S7 and S8 in Δ*ppk* mutants compared to wild-type controls (**Fig. 1B and Supplementary Table 1**). Overall, our work demonstrates that PPK is required for a timely response to MOPS-induced starvation.

### Lack of amino acids does not explain Δ*ppk* mutant phenotypes

We detected upregulation of many amino acid biosynthesis and binding enzymes in wild-type controls compared to Δ*ppk* mutants (33% of significantly differentially expressed proteins, **Supplementary Table 1**). We wondered if a lack of available amino acids could account for the phenotypic and proteomic differences observed in our experiments. In agreement with previous work of Kurdo et al. [8], we found that supplementation of MOPS media with 0.05 % amino acids improved growth of Δ*ppk* mutants (**Fig. 2A**). However, this increase in growth was not enough to match the growth rate of wild-type cells, which also increased in response to amino acid addition. At the protein level, addition of amino acids to MOPS media increased the expression of YbdL-3Flag, with minor increases for MetE-3Flag, YeaG-3Flag, and OtsA-3Flag in wild-type cells (**Fig. 2B**). However, in Δ*ppk* mutants, addition of amino acids failed to rescue protein expression to levels seen in untreated wild-type cells. Together, these data suggest that while amino acid deficiencies may contribute to some phenotypes of Δ*ppk* mutant cells, they are unlikely to explain the broad protein dysregulation observed in our proteomics dataset. Instead, we postulate that wild-type cells respond to MOPS-induced stress more promptly than Δ*ppk* cells by modulating the expression of multiple pathways. Investigation of these distinct pathways will uncover new insights into PPK and polyP modes of action.

**Figure 2.**
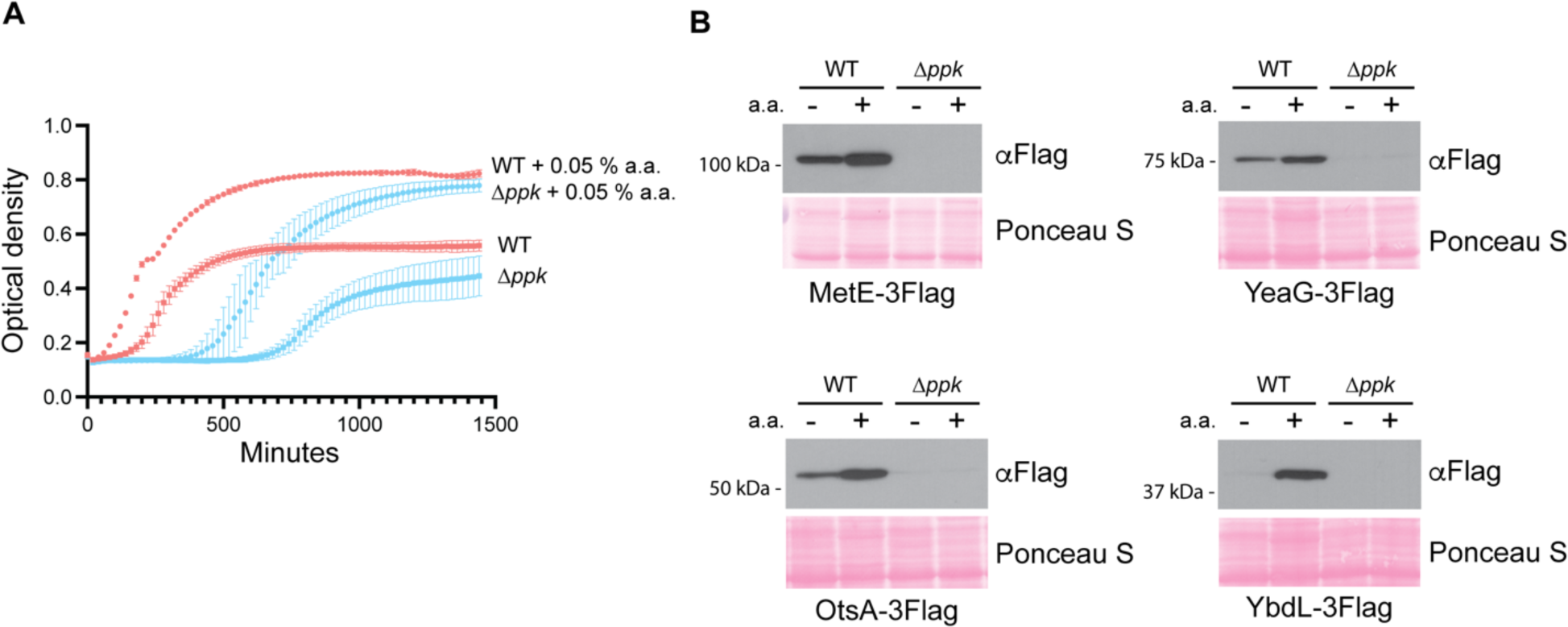
Impact of amino acid deficiencies on growth and proteome regulation in Δ*ppk* mutants. **A)** Growth of wild-type and Δ*ppk* mutant cells in MOPS media with and without 0.05% amino acid supplementation. Cells were grown in LB to mid-exponential phase and then diluted to 0.1 OD_600_ in MOPS minimal media in the absence or presence of 0.05% amino acids. Growth was monitored using the BioscreenC plate reader (37°C with shaking, wavelength 600 nm). Error bars represent standard deviation of the mean for 3 biological replicates. Error bars not shown for data points where standard deviation is smaller than size of the symbol itself. **B**) Effect of amino acid supplementation on expression of significantly differentially expressed proteins. Cells were grown to mid-exponential phase in LB and then shifted to MOPS minimal media in the presence or absence of 0.05% amino acids for 3 hours. Protein extracts were resolved using a 12% SDS-PAGE gel, transferred to PVDF membrane, and proteins detected using an anti-Flag antibody. Images are representative of results from ≥3 experiments.

### PPK plays a role in regulating expression of proteins required for lipid A modification

We were intrigued by the proteins ArnB, ArnC and EptA, which were only detected in wild-type control but not in Δ*ppk* mutant samples, because they function in pathways associated with cationic antibiotic resistance [27, 36–38]. The Arn proteins (ArnA, B, C and D) synthesize the donor substrate for the L-Ara4N modification, undecaprenyl-phosphate-L-Ara4N, on the cytoplasmic side of the inner membrane [37–39] (**Fig. 3A**). The undecaprenyl substrate is then flipped across the membrane by ArnE and ArnF [40] (**Fig. 3A**). The glycosyl transferase ArnT then transfers L-Ara4N from the undecaprenyl donor to the lipid A domain of the LPS at the periplasmic face of the inner membrane [41] (**Fig. 3A**). Like ArnT, the active site of EptA that is responsible for pEtN modification resides in the periplasm [42] (**Fig. 3A**). Lastly, both pEtN and L-Ara4N modified lipid A are transported to the outer membrane by the LPS transport system [43]. For more information on the pathway, see review by Whitfield and Trent [44].

**Figure 3.**
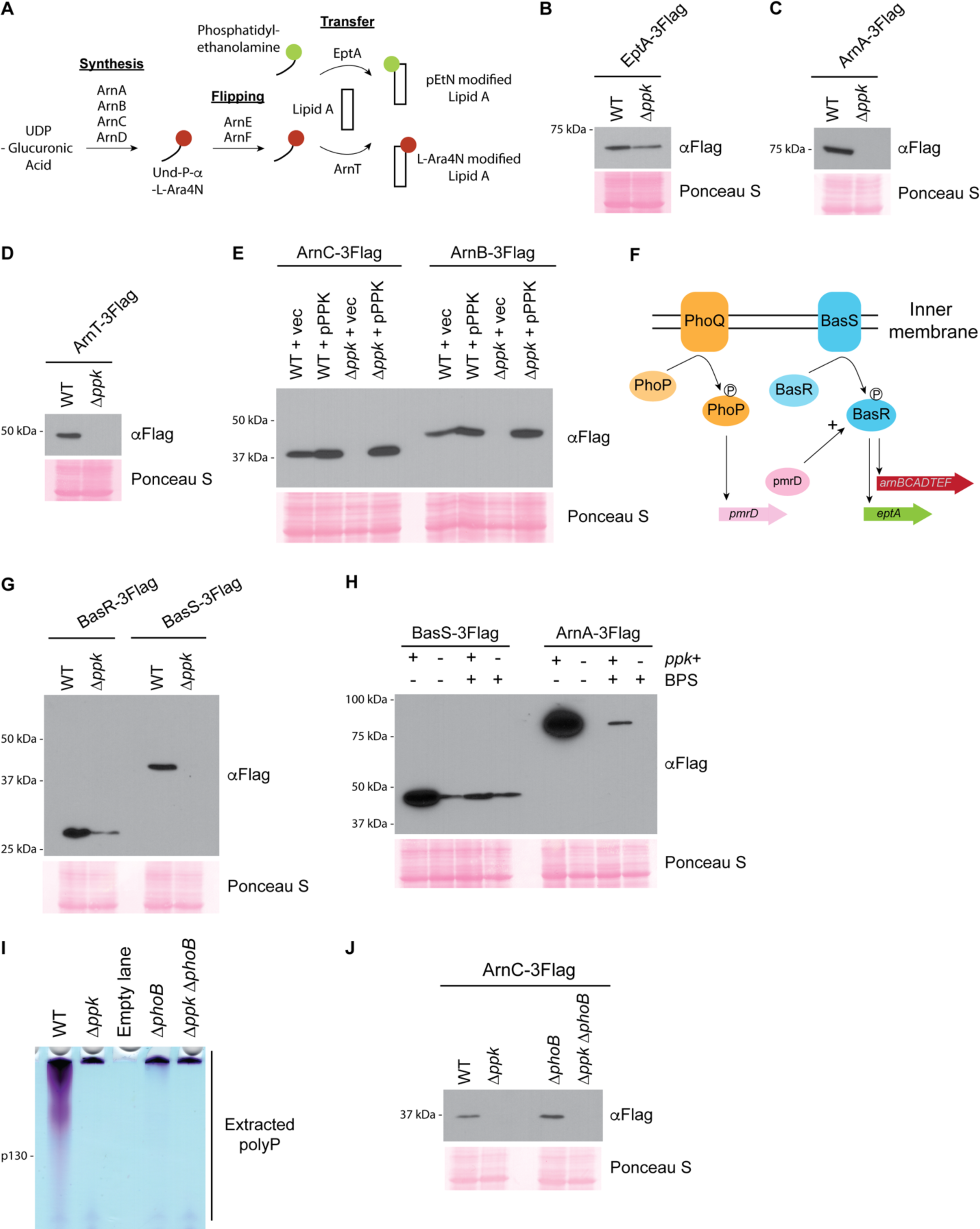
A) PPK positively regulates the BasS/R transcriptional circuit during starvation. **A)** Schematic for Arn and EptA-catalyzed lipid A modifications in *E. coli.* ArnA, ArnB, ArnC, ArnD synthesize the donor substrate (Und-P-Und-P-α-LAra4N). ArnE and ArnF flip and transport the donor substrate to ArnT. ArnT and EptA transfer their respective modifications to newly synthesized lipid A molecules. Note that pEtN and L-Ara4N are shown as single modifications, but doubly modified species containing 2 moieties (total) of pEtN and/or L-Ara4N are also possible. **B**)**-E**) Expression of EptA (B), ArnA, (C) and ArnT (D) and rescue of Arn expression (E) following MOPS starvation. Extracted protein samples were resolved using SDS-PAGE, transferred to PVDF and detected using an anti-Flag antibody. Note that polar effects due to tagging may influence *eptA* regulation and visualized expression in both wild-type and mutant strain backgrounds. **F)** Schematic showing induction of *arnBCADTEF* and *eptA* transcription by the PhoQ/PhoP and BasS/BasR two-component systems. **G)** BasS and BasR expression following MOPS starvation. Proteins were extracted from the indicated strains and analyzed as described above. **H)** Role of iron in BasS and ArnA expression. BPS was used to chelate iron from MOPS media, following the switch from LB. At the 3-hour timepoint, proteins were extracted and analyzed as described above. **I)** PolyP accumulation in Δ*phoB* mutants. Polyphosphate was extracted from the indicated strains grown in MOPS media and analyzed on a TBE-urea gel and stained with toluidine blue. **J)** Expression of ArnC-3Flag by Δ*phoB* mutants. Following MOPS starvation, proteins were extracted from the indicated strains and analyzed as described above. The same strains were used for 3I and 3J. Images shown are representative of results from ≥3 experiments, except for (I), which is representative of 2 independent replicates.

Together, L-Ara4N and pEtN modifications decrease the net negative charge of the outer membrane and reduce the interaction with cationic antimicrobial peptides such as polymyxin [45]. As we found for ArnC-3Flag and ArnB-3Flag in **Fig. 1C**, expression of EptA-3Flag was also PPK dependent (**Fig. 3B**). Other Arn proteins were not detected in our mass spectrometry experiment, but we confirmed the same trend for ArnA-3Flag and ArnT-3Flag (**Fig. 3C & Fig. 3D**). Importantly, Arn protein levels could be restored by plasmid-based expression of *ppk* from its endogenous promoter (pPPK, **Fig. 3E**). In fact, cells expressing pPPK in either wild-type or *Δppk* mutant backgrounds had somewhat higher levels of Arn proteins compared to wild-type strains with empty vector controls (**Fig. 3E**). We surmise that PPK is somewhat overexpressed in both strain backgrounds due to multiple copies of the plasmid and that Arn expression scales with total PPK levels.

Next, we asked if the switch from LB to MOPS media was triggering Arn expression. As anticipated, Arn proteins were low for both wild-type and Δ*ppk* mutants during growth in LB, and it was only during prolonged growth in MOPS that their levels increased in wild-type cells (**Supplementary Fig. 2A**). This led us to check if regulation depended on the BasS/BasR two-component system, which sits upstream of the *arnBCADTEF* operon and *eptA* gene, and responds to various stresses (**Fig. 3F**). In *E. coli,* the BasS membrane protein auto-phosphorylates in response to high concentrations of iron and zinc, and subsequently trans-phosphorylates the transcription factor BasR to promote transcription of *arnBCADTEF* and of *eptA,* which is found in the same operon as *basS* and *basR* [46–49]. In parallel, the PhoQ/PhoP two-component system, activated under conditions of low magnesium promotes BasR activation via PmrD (**Fig. 3F**) [50, 51]. We found that expression of both BasS and BasR were decreased in Δ*ppk* mutant cells, compared to wild-type cells, during starvation (**Fig. 3G**). In contrast, we found that PhoP and PhoQ protein levels remained largely unchanged (**Supplementary Fig. 2B & Supplementary Fig. 2C**). Thus, while we cannot rule out additional points of regulation, our data support a model wherein Arn and EptA expression during starvation in MOPS depends on PPK and/or polyP regulation of the BasS-BasR two-component system. There are several reports of BasR and BasS regulation by the stationary phase sigma factor RpoS [52, 53]. However, we found that Δ*ppk* mutants had RpoS protein levels that were similar to those of wild-type cells during starvation in MOPS (**Supplementary Fig. 2D**), suggesting additional modes of action.

To our knowledge this is the first description of *E. coli* EptA and Arn protein induction by MOPS media, and the mechanism at play is unknown. In LB media, the BasS/BasR transcriptional circuit induces Arn expression and downstream modifications in the presence of high iron levels (>200 µM) [46, 54], but the iron concentration in MOPS is quite low (10 µM). Still, we remained curious about the role of iron based on a previous report that MOPS-induced polyP can bind iron to inhibit the Fenton reaction, which decreases the production of reactive oxygen species [16]. Consistent with a requirement for iron in BasS/R induction, treatment with iron chelator BPS blunted expression of Arn proteins in wild-type cells grown in MOPS (**Fig. 3H**). However, we note that loss of *ppk* did not impact expression of Arn proteins in LB media treated with high iron (**Supplementary Fig. 2E**). Thus, while iron is important for BasS/R induction in both LB and MOPS, the impact of *Δppk* is unique to MOPS. We reasoned that BasS/R regulation by PPK could depend on polyP accumulation, which occurs in MOPS, but not in LB treated with iron (**Supplementary Fig. 2F**). To test this idea, we analyzed Arn protein induction in Δ*phoB* mutants, which are deficient in polyP accumulation even in MOPS media (**Fig. 3I)** [5, 28]. To our surprise, Δ*phoB* cells had levels of Arn expression comparable to wild-type controls (**Fig. 3J**), suggesting that additional PPK activities beyond polyP synthesis may contribute to the molecular phenotypes described here (See Discussion).

### pEtN and L-Ara4N modifications are downregulated in Δ*ppk* mutants

Next, we directly examined lipid A modifications that depend on Arn and EptA expression, namely L-Ara4N and pEtN addition. For these experiments, we used the W3110 strain background that has been used extensively for lipid A analyses [55, 56]. As a control, these strains showed PPK-dependent expression of Arn proteins in MOPS media, similar to what we observed in the MG1655 background used for our other assays (**Supplementary Fig. 3**). A time course of protein expression showed that in wild-type cells, ArnC-3Flag was detectable after 3 hours in MOPS media and remained upregulated for the duration of the experiment (**Fig. 4A**). In contrast, ArnC-3Flag was undetectable in Δ*ppk* mutants after 6 hours in MOPS media and was only observed after overnight growth (**Fig. 4A**). Therefore, we chose 6 hours (3 hours post-Arn upregulation) as a timepoint to measure lipid A modifications using radio-labelling and thin-layer chromatography. Indeed, compared to their wild-type counterparts, we saw that Δ*ppk* mutants were defective in the accumulation of lipid A species singly or doubly modified with pEtN and L-Ara4N modifications (**Fig. 4B**). This defect was fully rescued by introduction of the pPPK plasmid (**Fig. 4B**). Therefore, differences in levels of Arn and EptA proteins translate to changes in lipid A modification between WT and Δ*ppk* mutants.

**Figure 4.**
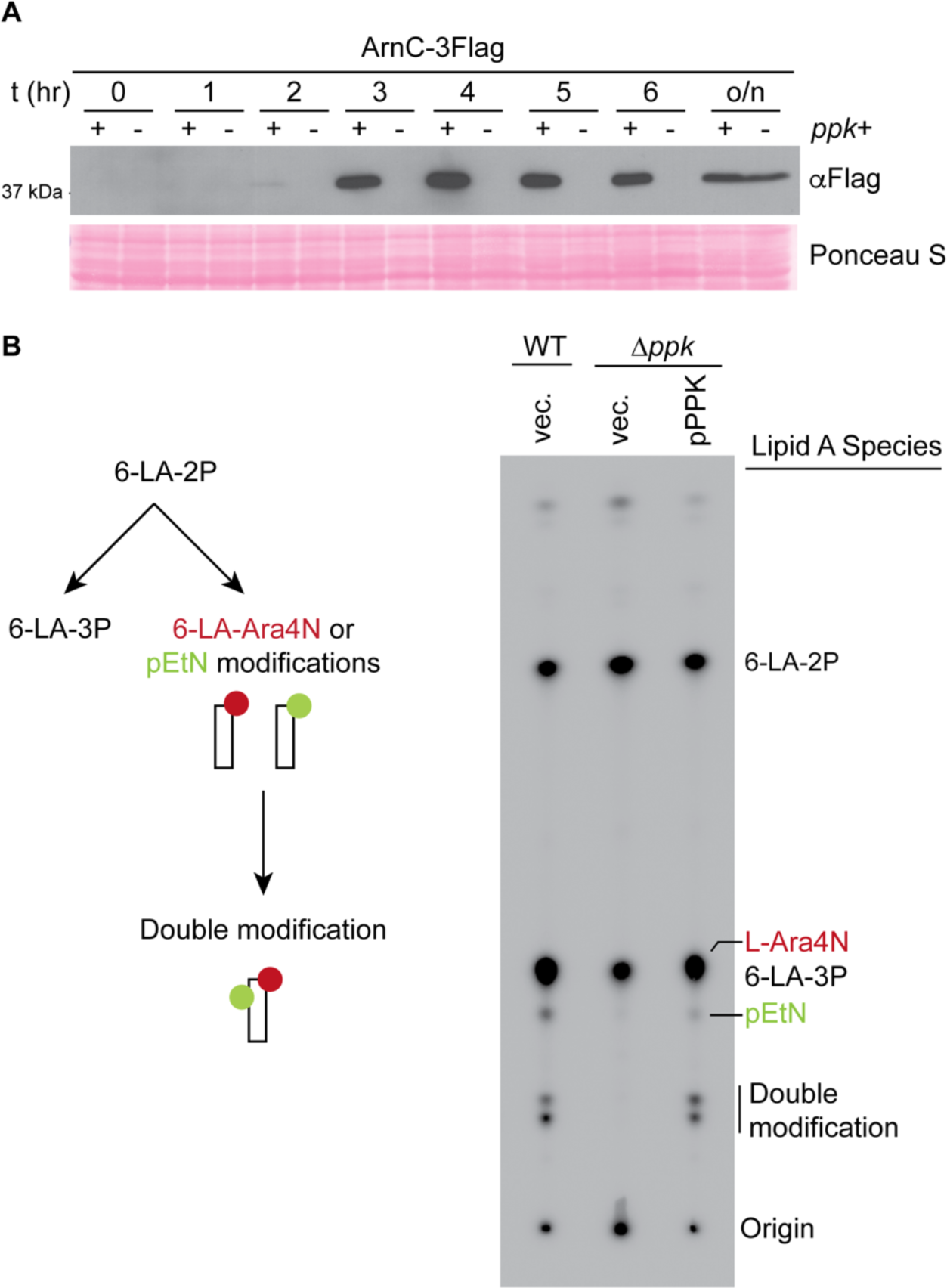
PPK promotes accumulation pEtN and L-Ara4N modifications on lipid A. **A)** Time course of ArnC expression following the shift from LB to MOPS media. Expression was analyzed for the indicated strains at the timepoints shown. Protein samples were resolved using a 12% SDS-PAGE gel, transferred to PVDF membrane, and proteins detected using an anti-Flag antibody. **B**) Schematic of lipid A modification (left) and lipid A profiles (right) of strains mutated for or expressing *ppk*. The indicated strains were grown in LB until mid-log phase before being shift to MOPS media supplemented with ^32^P for 6 hours. Lipid A was isolated as described in the Materials & Methods and analyzed via thin-layer chromatography prior to phosphorimaging. ^32^P-labelled lipid A species are labelled on the right side of the image. During growth in LB, the majority of the lipid A is hexa-acylated and *bis*-phosphorylated (6-LA-2P) and ∼1/3^rd^ is modified with an additional phosphate group (6-LA-3P). Upon stimulation of BasS/R induction, 6LA-2P is instead used as the substrate for EptA and ArnT to generate pEtN and L-Ara4N-modified lipid A species. Note that the singly modified 6-LA-Ara4N species is not resolved from the 6-LA-3P species. ‘Doubly-modified’ refers to lipid A species carrying two moieties (total) of pEtN and/or L-Ara4N. LA, lipid A; 6-LA-2P, Lipid A with 2 phosphate groups; 6-LA-3P, lipid A with 3 phosphate groups. Images shown are representative of results from ≥3 biological replicates.

### PPK promotes polymyxin resistance

Positively charged L-Ara4N and pEtN modifications play a role in resistance to cationic antimicrobial peptides that alter membrane permeability and structure [57–59]. To study the phenotypic consequences of disrupting PPK-dependent regulation of lipid A modifications, we focused on polymyxin antibiotics, used both topically to treat Gram-negative bacterial infections, and systemically as a last-resort antibiotic in the clinic [60]. For these experiments, we used a polymyxin resistant strain (WD101) that carries a constitutively active *basR* allele (*basR^C^*) resulting in lipid A that is heavily modified with L-Ara4N and pEtN [61]. This strain is otherwise isogenic to W3110 [61]. Importantly, compared to wild-type cells, Arn protein expression was decreased in Δ*ppk* mutants carrying this allele as observed previously in other backgrounds (**Supplementary Fig. 3**). Using dilution assays, we confirmed that the *basR^C^* strain is resistant to polymyxin compared to the wild-type W3110 counterparts grown on MOPS (**Fig. 5A**) and this resistance required ArnA (**Supplementary Fig 4**). Deletion of Δ*ppk* decreased polymyxin resistance in the *basR^C^* strain (**Fig. 5A**), and this effect could be reversed by introduction of the pPPK plasmid (**Fig. 5B**). These experiments demonstrate that *ppk* contributes to *basR*^C^-mediated polymyxin resistance under starvation conditions.

**Figure 5.**
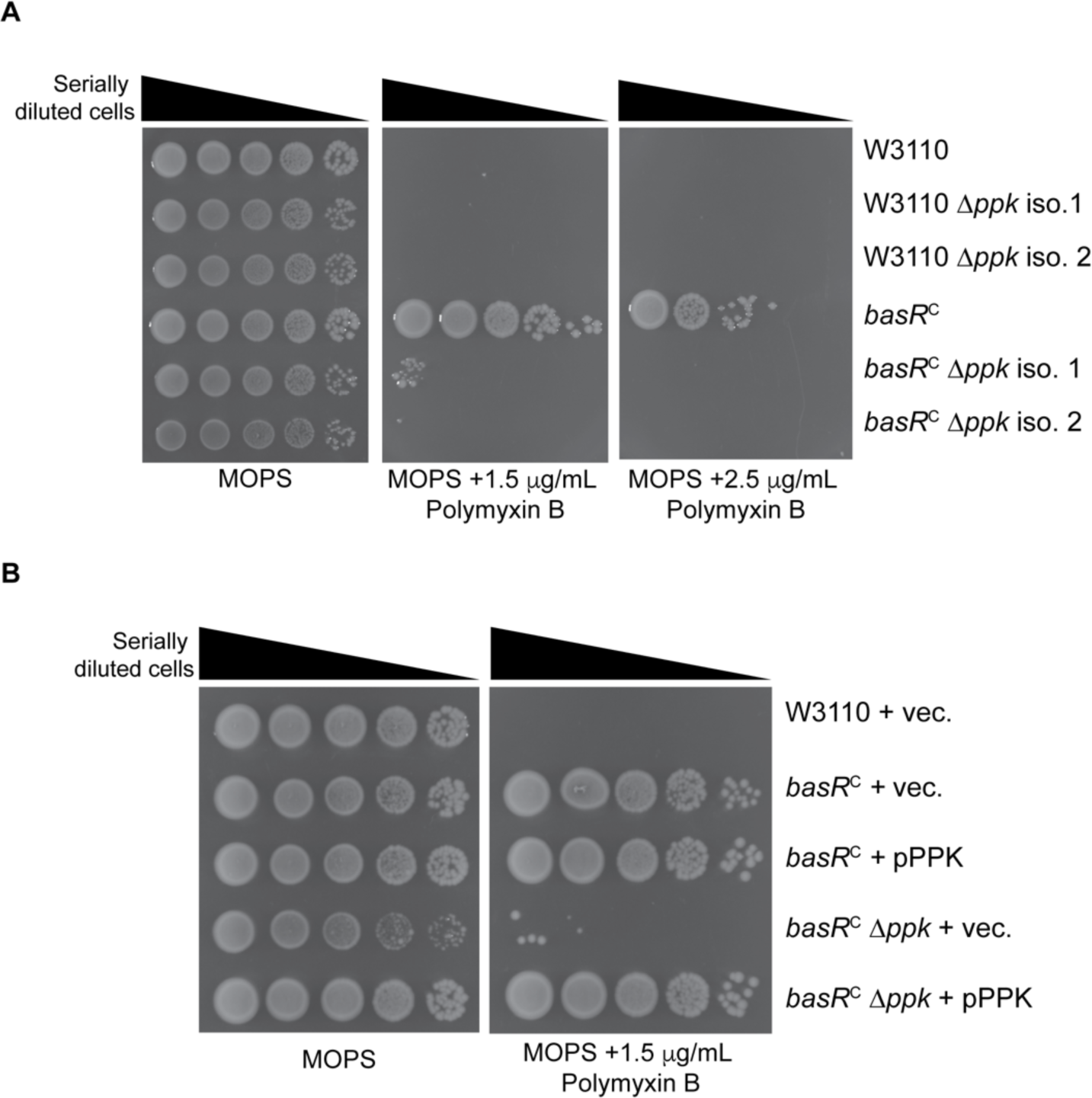
Polymyxin sensitivity of Δ*ppk* mutants. **A) & B)** Polymyxin growth phenotypes of Δ*ppk* mutants. The indicated strains were spotted in 10-fold serial dilutions on the indicated media and incubated at 37 °C for 2 days prior to imaging. Images shown are representative of results from ≥3 biological replicates.

## DISCUSSION

Our study is the first whole proteome analysis in *E. coli* comparing wild-type cells to Δ*ppk* mutants under conditions permissive for polyP synthesis. Our work will serve as a resource for researchers interested in how PPK and polyP control protein homeostasis, and, more broadly, how bacterial cells respond to starvation. We find that Δ*ppk* mutant cells remain poised for growth during starvation at the expense of upregulating biosynthetic pathways for nutritional building blocks such as amino acids. We further elaborate a critical role for PPK in regulating the conserved BasS-BasR two-component system and downstream expression of proteins involved in lipid A modifications. Finally, we uncovered evidence that PPK-dependent differences in lipid A modification contribute to polymyxin sensitivity. Together, our results illuminate a previously unexplored role of PPK in lipid A modification and antibiotic resistance.

The simplest interpretation of our data, illustrated in **Fig. 6**, is that PPK and/or polyP promote expression or activation of the BasS sensor protein and its cognate transcription factor BasR. In turn, BasR acts to stimulate transcription of both itself and genes encoding the EptA and Arn proteins. In support of the idea that the BasS/R-Arn circuit is regulated by polyP, as distinct from PPK, we only observed defects in Arn protein expression in MOPS media where polyP is synthesized to high levels. In LB media supplemented with iron, where polyP is absent, no differences in Arn protein expression was observed between wild-type strains and Δ*ppk* mutants. However, we note that a proposed role for polyP is complicated by the observation that Δ*phoB* strains express ArnC-3Flag protein at a level similar to wild-type cells, even though they are largely deficient in polyP accumulation. Based on this finding, it is possible that an unknown activity of PPK may underlie the molecular and cellular phenotypes described here. Alternatively, a non-zero level of polyP has been detected in Δ*phoB* strains previously [5], and this pool of polyP, or at least its cyclical synthesis and destruction, may be sufficient to drive BasS/R expression. In this scenario, polyP could interact directly with BasS-BasR or as-yet-unknown transcriptional regulators to promote transcription of the *basSbasR* operon.

**Figure 6.**
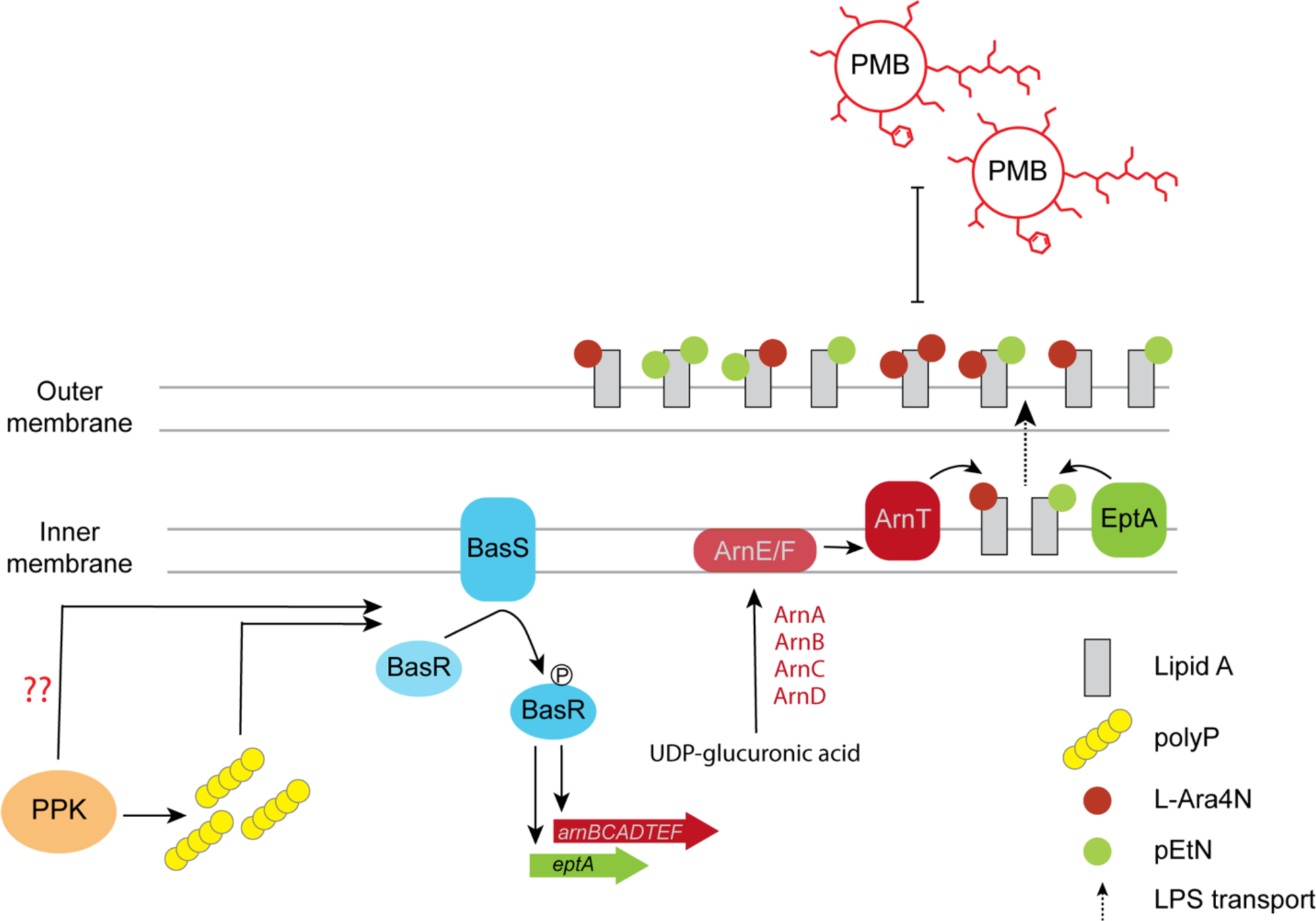
Role of PPK in the regulation of lipid A modification and polymyxin resistance. PolyP synthesized by PPK upon a switch from LB to MOPS media triggers BasS activation by autophosphorylation. Activated BasS then transphosphorylases BasR to induces downstream transcription of the *arnBCADTEF* operon and *EptA* gene. This results in increased level of Arn and EptA proteins, and upregulation of the respective L-Ara4N and pEtN modifications. Dashed arrows indicate an additional step where the modified lipid A (a key structural component of the LPS) is transported to the outer membrane by the LPS transport system. This reduces the net negative charge of the outer membrane and results in polymyxin B (PMB) resistance. What is still unknown is whether polyP is acting directly in BasS activation and if PPK has a role independent of polyP synthesis.

The impact of PPK could also be several steps removed from the transcriptional activities occurring directly at *basSbasR*. Here, it is particularly relevant that the trigger for activation of the BasS/R transcriptional program during MOPS starvation is unknown. The failure of wild-type cells to upregulate Arn expression in the presence of the iron chelator BPS suggests an iron-dependent response. However, given that induction of Arn protein expression occurs abruptly after three hours, we theorize that change in some other component of the media is also required. Cells lacking *ppk* may reach this point more slowly, due either to defects in a specific metabolic pathway, or as a result of overall slow growth in MOPS media. If true, we have uncovered an important consequence of Δ*ppk* cells failing to adapt to starvation conditions. Ultimately, finding the trigger for BasS/R activation in MOPS will provide important clues to unknown functions of PPK and new insights into the regulation of lipid A metabolism.

Collectively, our findings underscore that the pathways regulating modification of *E. coli* LPS, and the associated changes in antibiotic resistance, are intimately coupled to nutrient availability. In the context of host infection, the validity of PPK as a target to sensitize bacteria to cationic peptides like polymyxin may depend on whether bacteria at the site of infection are exposed to conditions that activate PPK. Intriguingly, there is evidence to suggest that bacteria colonizing in the center of dense biofilms experience starvation concomitant with decreased susceptibility to antibiotics [62–64]. We theorize that PPK may play an important role in this context. Notably, transcription of the *ppk-ppx* regulon does not change significantly during MOPS treatment [29] and the immediate molecular events that activate PPK during stresses of any kind remain unclear [29, 65–67]. Our work predicts that genes encoding PPK activators are also likely to play a role in polymyxin resistance.

In general, the players involved in LPS modifications, including the pEtN and L-Ara4N modifications, are highly conserved across Gram-negative bacteria [57]. Yet, there are important differences between some species in the details of regulation, such as in the crosstalk between two-component systems [50, 68]. As such, it is imperative to test if PPK and polyP impact these modifications in other bacteria. We anticipate that comparing and contrasting the role of PPK across multiple species will help to elucidate molecular mechanisms governing antibiotic sensitivity and the acquisition of resistance.

## MATERIALS AND METHODS

### Bacterial strains, media, and growth

All bacterial strains and plasmids, as well as their sources, used in this work are listed in **Supplemental Table 3**. Reagents are listed in **Supplemental Table 4**.

Tagged and deletion strains were generated using lambda-red mediated site-specific recombination via the arabinose or heat shock inducible systems expressed from pKD46 [69] and pSIM6 [70] plasmids, respectively. Endogenous kanamycin deletion and C-terminal 3Flag-kanamycin tagging cassettes were amplified from pKD4 [69] and pSUB11 [71], respectively. In select cases (**Supplemental Table 3**), resistance markers were excised using FLP recombinase activity expressed from pCP20 [72]. Antibiotics were added when appropriate: kanamycin (50 µg/mL), ampicillin (100 µg/mL). For recombineering genetic transformations cells were made electrocompetent and transformed using protocols described previously [73]. Plasmids were also transformed by electroporation.

#### (1) Nutrient downshift

Strains were grown in LB media at 37°C unless carrying pSIM6 and pKD46 plasmids which were grown at 30 °C. For nutrient downshift experiments overnight cultures, grown in LB media, were diluted to 0.1 OD_600_ in LB the next day and grown to mid-exponential phase (∼0.6 OD_600_). Cells were then washed twice with 1xPBS and resuspended in MOPS minimal media (Teknova^TM^) supplemented with 0.1 mM K_2_HPO_4_, 0.4% glucose. Where indicated, casamino acids (Bacto^TM^) were used for amino acid supplementation at a final concentration of 0.05%. Cells were grown in minimal media for 3 hours before harvesting on ice. Cell pellets were flash frozen using dry ice and stored at −80°C.

#### (2) Other growth conditions

Iron in MOPS media was chelated using 0.2 mM bathophenanthrolinedisulfonic acid disodium salt hydrate (BPS). BPS was added to MOPS media at the start of the nutrient downshift. Where indicated, LB media was supplemented with 0.2 mM iron sulfate when overnight cultures were diluted to 0.1 OD_600_. LB minus and plus iron cultures were grown to mid-exponential phase (about 1.5 hours) before harvesting cells.

### Mass spectrometry

#### (1) Cell growth

Five overnight cultures (n=5) were prepared for each wild-type and *ppk* strains from freshly streaked plates. The next day, overnight cultures were diluted in 100 mL LB and grown to mid-exponential phase, washed two times with 1xPBS and then switched to MOPS minimal media without amino acids for three hours as described above (methods: nutrient downshift). 60 OD_600_ and 5 OD_600_ equivalents of cell volume were harvested for mass spectrometry analysis and polyP extraction, respectively.

#### (2) Protein extraction and precipitation

Protein extracts were prepared by trichloro-acetic acid (TCA) precipitation. Cells pellets were thawed on ice and resuspended in 700 µL mass spectrometry-grade lysis buffer (5%SDS, 100 mM tetraethylammonium bromide (TEAB), Roche cOmplete protease inhibitor cocktail tablets). The cell suspension was lysed by sonication on ice for 30 seconds at power level 3 with 1 minute break in between, for a total of 3 times. The cell lysate was then centrifuged at 13 000 rpm, 4°C for 15 minutes. The supernatant was transferred to a new tube and centrifuged again for 10 minutes prior to being collected.

Proteins were precipitated by adding 2.8 mL 15% TCA in acetone to the cell lysate (brings final concentration of TCA to 10%), mixing by inverting and incubating at −20°C overnight. The next day, the samples were centrifuged at max speed, 4°C for 10 minutes and supernatant was discarded immediately after the spin. The pellet was air dried for 30 mins, then resuspended in dissolution buffer (5% SDS, 100 mM tetraethylammonium bromide) prior to BCA quantification and lyophilization. Lyophilized samples were sent to UC Davis Proteomics Core for mass spectrometry analysis.

#### (3) Sample preparation

The following protocols (3-5) were provided by the UC Davis Proteomics Core with minor alterations.

Lyophilised proteins were solubilised in 50 µL of solubilization buffer, consisting of 5% SDS, 50 mM triethyl ammonium bicarbonate, pH 7.5. A Bichinoic Acid Assay was taken of all samples, and 100 µg total protein from each sample was then used to perform protein digestion via suspension-trap devices (S-Trap) (ProtiFi). Disulfide bonds were reduced with dithiothreitol and alkylated with iodoacetamide in 50mM TEAB buffer. The enzymatic digestion consisted of an addition of trypsin at 1:100 enzyme: protein (wt/wt) for 4 hours at 37 °C, followed by a boost addition of trypsin using same wt/wt ratios for overnight digestion at 37 °C. Peptides were then eluted from the S-Trap by sequential application of elution buffers of 100 mM TEAB, 0.5% formic acid, and 50% acetonitrile 0.1% formic acid. The eluted tryptic peptides were dried in a vacuum centrifuge prior to re-constitution in 0.1% trifluoroacetic acid. These were subjected to Liquid Chromatography couple to tandem Mass Spectrometry (LC-MS/MS) analysis as described below.

#### (4) Liquid Chromatography

Peptides were resolved on a Thermo Scientific Dionex UltiMate 3000 RSLC system using a PepMap 75 µm x 25cm C18 column with 2 μm particle size (100 Å pores), heated to 40 °C. A final volume of 5 μL was injected, corresponding to 1 μg of total peptide, and separation was performed in a total run time of 90 min with a flow rate of 200 μL/min with mobile phases A: water/0.1% formic acid, and B: 80%ACN/0.1% formic acid. Gradient elution was performed from 10% to 8% B over 3 min, from 8% to 46% B over 66 min, and from 46 to 99% B over 3 min, and after holding at 99% B for 2 min, down to 2% B in 0.5 min followed by equilibration for 15min.

#### (5) Mass spectrometry

The peptides were analyzed on an Orbitrap Fusion Lumos (Thermo Fisher Scientific) mass spectrometer. Spray voltage was set to 1.8 kV, RF lens level was set at 46%, ion transfer tube temperature was set to 275 °C. The mass spectrometer was operated in a data-dependent acquisition mode. A survey full scan mass spectra (from m/z 375 to 1600) was acquired in the Orbitrap at a resolution of 60,000 (at 200 m/z). The automatic gain control (AGC) target for MS1 was set as 4e5, and ion filling time was set as 50 msec. The n=15 most abundant precursor ions with charge state +2, +3 were isolated in a 3-sec cycle, isolation window width of 1.2 m/z, fragmented by using collision-induced dissociation (CID) fragmentation with 30% normalized collision energy, and detected via IonTrap, with a scan rate set to Rapid. The AGC target for MS/MS was set as 5e3 and ion filling time was set at 35 msec. The dynamic exclusion was set to 50 sec with a 10-ppm (parts per million) mass window.

### Bioinformatics analysis

#### (1) Protein identification

Mass spectrometry RAW data were processed using the Trans-Proteomic Pipeline (TPP v5.2.0) [74]. Files from the mass spectrometry runs were converted to mzML files using the msconvert tool from ProteoWizard [75] (v3.0.22088). Comet [76] (v2018.01.04) was used to search the files against the UniProt [77] *E. coli* protein sequence database (UP000000625 downloaded 2021-08-04), along with a target-decoy strategy where all protein sequences were reversed. The database search was performed with trypsin as a digestive enzyme, allowing for up to 3 missed cleavages and considering semi-tryptic digestion. The peptide mass tolerance was set to 20 ppm. Carbamidomethylation of cysteine was set as a fixed modification, and the variable modifications considered were deamidation of asparagine and glutamine, as well as oxidation of methionine. The probability of protein identifications was evaluated with ProteinProphet [78], and proteins identified at a false discovery rate (FDR) < 1% were deemed confidently identified.

#### (2) Differential expression analysis

The spectral counts from confidently identified proteins were used for downstream analysis. Proteins with zero spectral counts in a given experiment were imputed a quantification value by random sampling of the lowest 20% of non-zero spectral counts in the entire dataset to account for missing values. All spectral counts were normalized by the total number of spectral counts in a given experiment. Differential expression was assessed on the normalized spectral counts using a two-tailed, two-sample Students *t*-test, assuming unequal variance. A *t*-test was performed for proteins that had at least 3 non-zero spectral counts out of 5 replicates prior to imputation in both experimental conditions, and the Benjamini-Hochberg [79] procedure was used to adjust *p*-values for multiple hypothesis testing. Proteins with an FDR-adjusted *p*-value < 0.05 were considered significantly differentially expressed. Finally, proteins that were identified in all replicates of one condition and not identified in any replicates of the other condition did not undergo a *t*-test, but still reflect a significant expression difference. As such, these were named “all-or-none” proteins and are reported as having differential expression.

#### (3) Gene Ontology enrichment analysis

Ontologizer [80] (v2.0) was used to identify Gene Ontology [33] annotations that were significantly enriched in the set of differently expressed proteins (FDR-adjusted *p*-value < 0.05) and “all-or-none” proteins. The enrichment was performed against a background of all proteins identified with mass spectrometry (confidently identified at an FDR < 1%). The OBO ontology file and the GAF annotation file used in the analysis were downloaded from http://geneontology.org/ on 2023-01-02. The *p*-values were adjusted using the Benjamini-Hochberg procedure, and Gene Ontology terms that were enriched with an adjusted *p*-value < 0.05 were considered significantly enriched.

#### (4) Gene set enrichment analysis

A Gene Set Enrichment Analysis (GSEA) [34] (v4.1.0) was performed to identify sets of Gene Ontology terms that were enriched in the entire set of proteins (confidently identified at an FDR < 1%). Gene sets were built of Gene Ontology terms and their annotated proteins, and GSEA excluded Gene Ontology terms that annotated more than 1000 proteins or less than 3 proteins to remove general and highly specific terms. 1000 gene set permutations were used to estimate enrichment scores. The enrichment scores were normalized with the “meandiv” parameter to allow for a more accurate comparison of enrichment scores across gene sets. Gene sets with a *q*-value < 0.05 were considered significantly enriched.

### Polyphosphate extraction

#### (1) Polyphosphate extraction

Cells were grown as indicated and polyP extraction was conducted as described previously [81]. For clarity, similar language is used to describe the protocol here. Five OD_600_ equivalents of pelleted cells were thawed on ice, resuspended in 400 µL of LETS buffer (100 mM LiCl, 10 mM EDTA, 10 mM Tris-HCl, 0.2% SDS) at 4°C, then transferred to a tube containing 600 µL of room-temperature neutral phenol (pH 8) and 150 µL of RNase-free water. Tubes were vortexed for 20 seconds and 600 µL chloroform was added. Next, tubes were again vortexed for 20 seconds, and then centrifuged for 2 minutes at 13,000 g. The top 600 µL layer was transferred to a new tube containing 600 µL of chloroform before vortexing for 20 seconds and centrifuging for 2 minutes at 13,000 g. The top 400 µL layer was transferred to a new tube and treated with 2 µL of 10 mg/mL RNaseA and DNaseI, each, for 1 hour at 37°C. Next, the mixture was transferred to prechilled tubes containing 1 mL 100% ethanol and 120 mM sodium acetate (pH 5.3) and left overnight at - 20°C to precipitate. The next day, samples were centrifuged for 20 min at 13,000 g for 20 mins at 4°C. Supernatant was discarded and 500 µL 70% ethanol was added before centrifuging for 5 min at 13,000 g at 4°C. Supernatant was again discarded and pellet was air dried to remove trace ethanol. The translucent polyP pellet was resuspended in 30 µL RNase-free water and stored at - 80°C.

#### (2) Gel analysis

Extracted polyP was visualized using a 15.8% TBE-urea gel (5.25 g urea, 7.9 mL 30% acrylamide, 3 mL 5xTBE, 150 µL 10% APS and 15 µL TEMED). Extracted polyP was mixed at a 1:1 ratio with loading dye (10 mM Tris-HCL pH 7, 1 mM EDTA, 30% glycerol and bromophenol blue) and 10 µL was loaded into the gel. Gels were run at 100 V for 1 hour and 45 minutes in 1xTBE as the running buffer. Three microliters of RegenTiss polyP standards p14 (20 mM), p60 (6.5 mM) and p130 (2.5 mM) were used. The gel was stained in fixing solution containing toluidine blue (25% methanol, 5% glycerol and 0.05% toluidine blue) for 15 minutes and washed several times with destaining solution (fixing solution without toluidine blue) before being left overnight to fully destain.

### Western blotting

Cells were grown as indicated in figure legends and described in the “Bacterial strains and growth conditions” section. Cell pellets (3 OD_600_ equivalent of cells) frozen at −80°C were thawed on ice and resuspended in 100 µL sample buffer (800 µL sample buffer 100 µL 1 M DTT, 100 µL 1.5 M Tris-HCl pH 8.8). Samples were boiled at 100°C for 10 mins, then centrifuged at 13,000 rpm for 2 mins and the supernatant was transferred to new tubes. Only ArnT and EptA membrane protein samples were prepared without boiling. Cell pellets resuspended in 100 µL sample buffer were sonicated at power level 1 for 12 seconds, then centrifuged at 13,000 rpm for 2 mins and the supernatant was transferred to new tubes. Protein samples were loaded on the indicated % of SDS-acrylamide gel. Proteins were transferred to PVDF membrane. Membranes were blocked for 20 minutes with shaking using TBST with 5 % milk and washed 3 times for 10 minutes each with TBST after incubation with the primary and secondary antibodies. Blots were exposed to autoradiography film from Thomas Scientific. Conditions for antibody use can be found in **Supplemental Table 4.** Scanned images were opened in Photoshop. In most cases small linear brightness and contrast adjustments were made to lighten the image background. Adjustments were applied evenly across the entire image shown.

### Liquid growth curves

Three biological replicates of each strain were cultured overnight in LB media. Overnights were diluted to 0.1 OD_600_ in LB and grown to mid-exponential phase (∼0.6 OD_600_). Cells (0.5 OD_600_ equivalent) were then washed twice with 1xPBS and resuspended in 50 µL of MOPS media (1xMOPS, 0.01 mM K_2_HPO_4_) without amino acids. Twenty-five microliters of the cell suspension was transferred to 1 mL of MOPS without and with 0.05% amino acids (final OD = 0.1 OD_600_). Two hundred microliters of cells were pipetted in technical replicates of 3 and growth was monitored using the BioscreenC plate reader set at 37 °C with continuous shaking. Optical density measurements were collected at a wavelength of 600 nm every 30 mins, 5 seconds after shaking stopped, for 24 hours.

### Polymyxin sensitivity

Indicated strains were streaked on LB or LB-kanamycin (vector carrying strains) plates and grown overnight at 37°C. The next day, a single colony from each strain was resuspended in 100 µL sterile water and serially diluted 10-fold in sterile water. Five microliter of each dilution was spotted onto the indicated plates that were prepared fresh on the day of use. Plates were allowed to dry prior to incubation at 37°C for 2 days before imaging. Linear brightness and contrast adjustments made in Photoshop were made evenly across the entire image shown.

### Lipid analysis

Overnight cultures were diluted 1:50 in fresh LB medium. Cells were harvested at OD_600_ of 0.6 and washed with 1X PBS. Bacteria were resuspended in 1X MOPS medium supplemented with 0.4% glucose, 0.1 mM K2HPO4 and 2.5 µCi/ml ^32^Pi. Labeled cells were grown at 37°C and harvested after 6h. Pellets were washed with 1X PBS. ^32^P-lipid A was extracted using mild acid hydrolysis as previously described [82]. The lipid A profiles were then resolved by thin-layer chromatography (TLC) in a solvent system consisting of chloroform, pyridine, 88% formic acid and water (50:50:16.5:5, vol/vol, respectively). The plates were exposed to a phosphor screen, and the radiolabeled lipids were visualized using an Amershan Typhoon phosphorimager system. Linear brightness and contrast adjustments were made in Photoshop to make clear the pEtN and L-Ara4N modifications. Linear adjustments were made evenly across the entire image shown.

## ACKNOWLEDGEMENTS

We thank members of the Downey Lab for critical evaluation of the manuscript. We thank the UC Davis Proteomics Core, especially Dr. Gabriela Grigorean for providing method details. We would also like to thank Dr. Michael Gray for the WT and Δ*ppk* parent strains, and pPPK plasmid. Lastly, we thank Dr. Lionello Bossi and Dr. Donald Court for the tagging and recombineering helper plasmids used in this work.

## Author contributions

Conceptualization: K.B., M.D. Methodology: K.B., I.A., C.M.H. Investigation: K.B., I.A., C.M.H. Supervision: S.M.T., M.L.A., M.D. Writing – original draft: K.B., M.D. Writing – review and editing: All authors.

## Data and materials availability

All data needed to evaluate the conclusions in this paper are present in the paper and/or the Supplementary Materials. Bacterial strains and plasmids used in this work can be provided by the Downey laboratory. The mass spectrometry proteomics data have been deposited to the ProteomeXchange Consortium [83] via the PRIDE [84] partner repository with the dataset identifier PXD043566. Reviewers can access the data with the following credentials: Username; Password: T0ib8hTL.

---

**Fig. S1.**
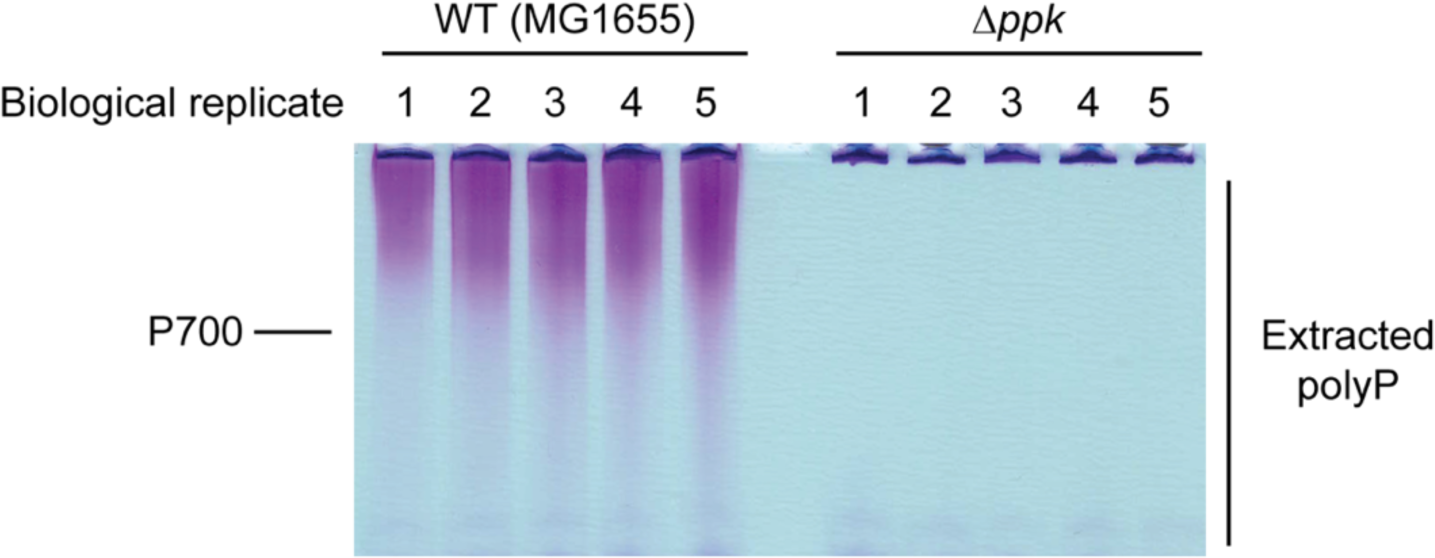
Wild-type *E. coli* accumulated polyP in MOPS minimal media while Δ*ppk* mutants do not. PolyP extraction gel from wild-type and Δ*ppk* mutant cultures used for mass spectrometry sample preparation. Overnight cultures were grown in LB media to mid-exponential phase and then shifted into MOPS minimal media for 3 hours to induce starvation and polyP accumulation. PolyP extracts were run on a TBE-urea gel and stained with toluidine blue. The migration of a chain ∼700 phosphate residues in length (p700) is indicated.

**Fig. S2.**
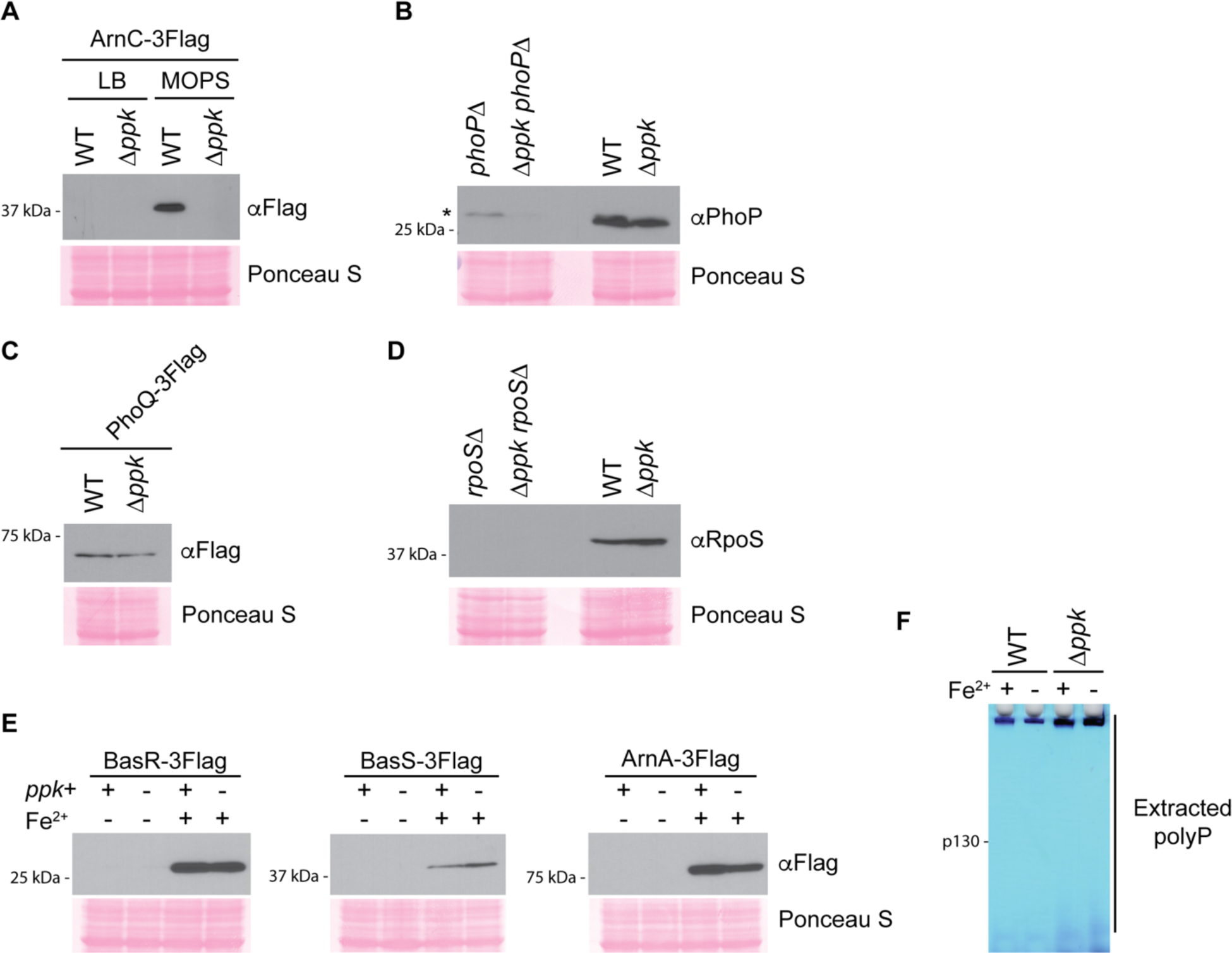
Molecular control of Arn and EptA protein expression by PPK. **A)** Induction of ArnC-3Flag expression upon the switch from LB to MOPS media. The indicated strains were grown in LB media to mid-log phase and shifted to MOPS media for 3 hours. Proteins were extracted and analyzed via SDS-PAGE prior to transfer to ta PVDF membrane and detection of tagged proteins with an anti-Flag antibody. **B)** Expression of PhoP between wild-type cells and Δ*ppk* mutants. The indicated strains were starved in MOPS media for 3 hours prior to protein extraction, separation by SDS-PAGE and detection with an antibody against PhoP. A background band in Δ*phoP* mutants is PPK-regulated, which makes evaluation of changes to PhoP protein expression difficult. Regardless, regulation of PhoP by PPK appears to be minimal. **C)** Expression of PhoQ-3Flag between wild-type cells and Δ*ppk* mutants. The indicated strains were starved in MOPS media and proteins were analyzed as described in (A). **D)** Expression of RpoS between wild-type cells and Δ*ppk* mutants. The indicated strains were starved in MOPS media for 3 hours prior to protein extraction, separation by SDS-PAGE, and detection with an antibody directed against RpoS. The untagged strain serves to validate the antibody. **E)** Expression of BasS-, BasR- and ArnA-3Flag in LB+iron by wild-type and Δ*ppk* mutants. The indicated strains grown in LB or LB + iron (200 µM FeSO_4_) for 1.5 hours prior to protein extraction, separation by SDS-PAGE, transfer to PVDF, and detection of tagged proteins with an anti-Flag antibody. **F)** Influence of iron on polyP accumulation. Polyphosphate was extracted from the indicated strains grown in LB or LB + iron (200 µM FeSO_4_) for 1.5 hours and analyzed on TBE-urea gels stained with toluidine blue. The same BasR tagged strains were used for 2E and 2F. Images shown are representative of results from ≥3 experiments, except for (F), which is representative of 2 independent replicates.

**Fig. S3.**
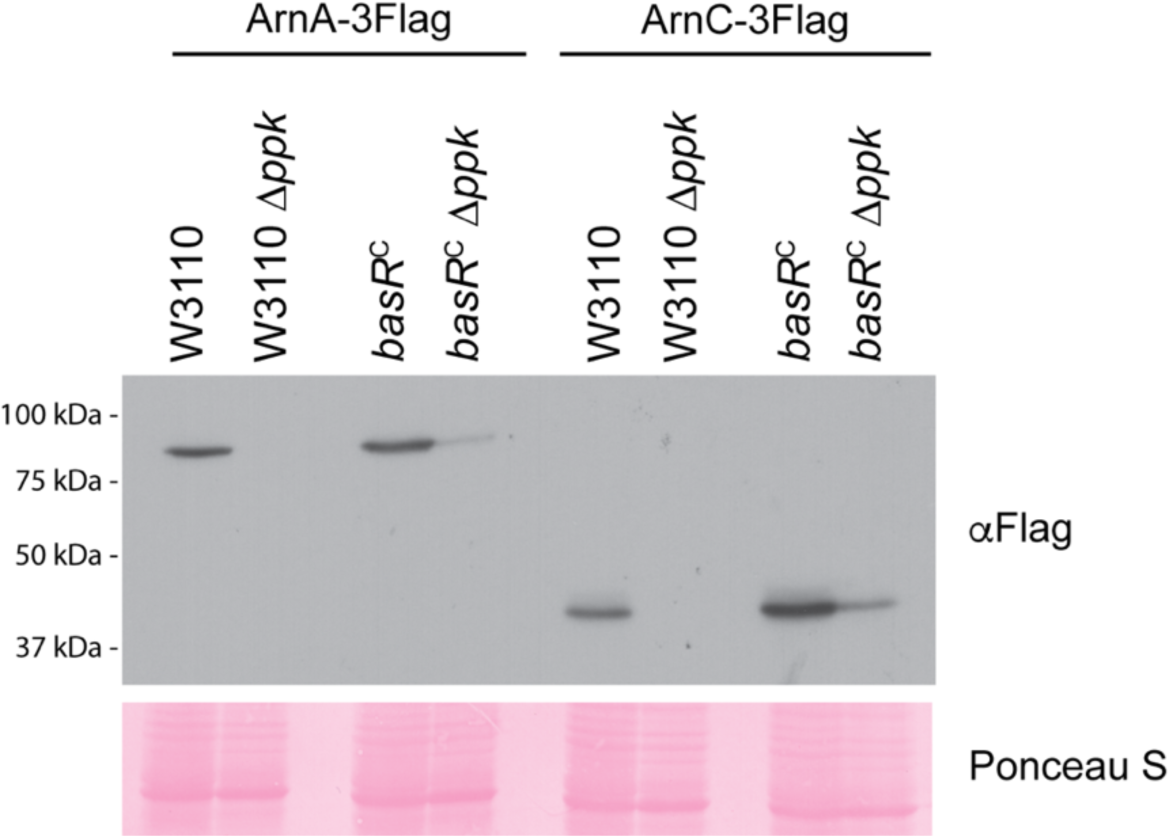
Regulation of Arn protein expression by PPK in the W3110 and WD101 (*basR^C^*) backgrounds. The indicated strains were grown in LB to mid log phase prior to shifting to MOPS for 3 hours. Proteins were extracted and separated via SDS-PAGE prior to transfer to PVDF membrane and detection with anti-Flag antibody. Images shown are representative of results from ≥3 experiments.

**Fig. S4.**
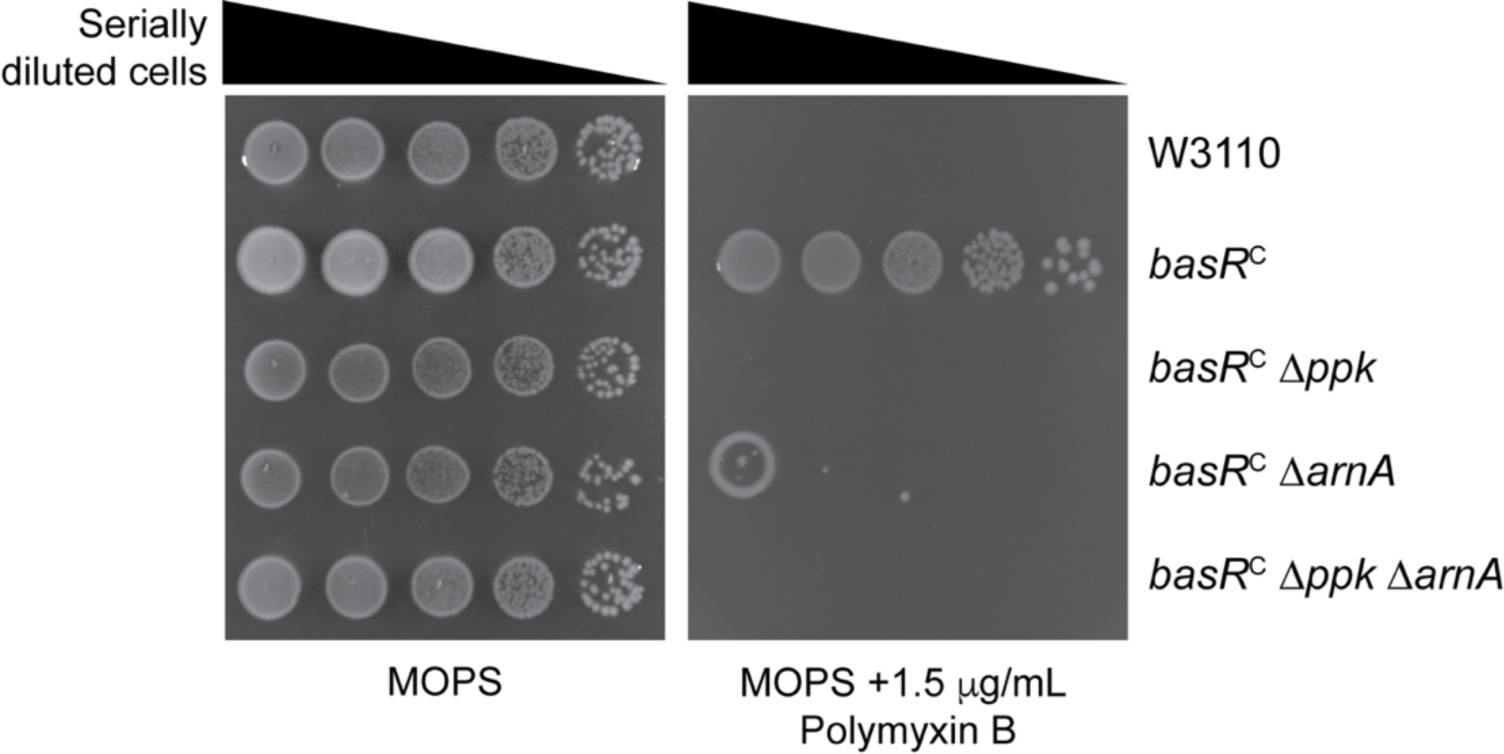
Arn-dependence of polymyxin resistance in *basR^C^* strains. The indicated strains were spotted in 10-fold serial dilutions on the indicated media and incubated at 37 °C for 2 days prior to imaging. Images shown are representative of results from ≥3 experiments.

**Table S1. (Attached as a separate file)**

Mass spectrometry data set.

**Table S2. (Attached as a separate file)**

Comparison of overlap between Varas *et al.* and Baijal *et al.* mass spectrometry data sets.

**Table S3. (Attached as a separate file)**

Bacterial strains and plasmids used for this work.

**Table S4. (Attached as a separate file)**

Reagents and antibody conditions used for this work.

